# Quantitative trait locus mapping for common scab resistance in a tetraploid potato full-sib population

**DOI:** 10.1101/2020.10.24.353557

**Authors:** Guilherme da Silva Pereira, Marcelo Mollinari, Xinshun Qu, Christian Thill, Zhao-Bang Zeng, Kathleen Haynes, G. Craig Yencho

## Abstract

Despite the negative impact of common scab (*Streptomyces* spp.) to the potato industry, little is known about the genetic architecture of resistance to this bacterial disease in the crop. We evaluated a mapping population (~150 full-sibs) derived from a cross between two tetraploid potatoes (‘Atlantic’ × B1829-5) in three environments (MN11, PA11, ME12) under natural common scab pressure. Three measures to common scab reaction were assessed, namely percentage of scabby tubers, and disease area and lesion indices, which were highly correlated (>0.76). Due to large environmental effect, heritability values were zero for all three traits in MN11, but moderate to high in PA11 and ME12 (0.44~0.79). We identified a single quantitative trait locus (QTL) for lesion index in PA11, ME12 and joint analyses on linkage group 3, explaining 22~30% of the total variation. The identification of QTL haplotypes and candidate genes contributing to disease resistance can support genomics-assisted breeding approaches.

## 1 Introduction

Common scab of potato (*Solanum tuberosum* L.) is an economically important disease that occurs worldwide. It is caused by pathogenic soil-borne bacteria belonging to several species in the genus *Streptomyces* (Loria et al., 1995; Wanner, 2009). Common scab is characterized by brownish superficial, raised or pitted lesions on tuber surfaces as a consequence of the phytotoxin thaxtomin A produced by pathogenic *Streptomyces* spp. (Loria et al., 1995; Kinkel et al., 1998). This disease is highly influenced by the environment (Haynes et al., 2010), especially by the soil conditions (Krištůfek et al., 2015), and by the virulence of the pathogens present in the soil (Wanner, 2006; Wanner and Haynes, 2009).

Although some management practices have been proposed to mitigate common scab damages (Dees and Wanner, 2012), there are currently no chemical or cultural management approaches that provide effective control of the disease. Breeding for varietal resistance is still being pursued as a more effective solution (Navarro et al., 2015; Braun et al., 2017b). However, the development of resistant varieties, as well as the study of the genetic architecture of resistance in potato, is quite challenging. Despite scab symptoms exhibiting a quantitative distribution (Haynes et al., 1997, 2009), scab resistance has been postulated as of oligogenic nature, with a dominant and a recessive locus (Murphy et al., 1995). Due to the autotetraploidy of potato (2*n* = 4*x* = 48), such a recessive locus would have to appear as in a quadruplex recessive state, which is not easy to achieve using conventional breeding (Bradshaw, 2017). In this sense, marker-assisted selection (MAS) could facilitate recessive allele introgression (Bethke et al., 2019).

Recent developments in genomics and bioinformatics tools have allowed most technical difficulties to be overcome when studying the genetic architecture of a trait in autopolyploid crops. For potato, strategies based on genotyping-by-sequencing (Uitdewilligen et al., 2013; Sverrisdóttir et al., 2017) or chip arrays (Felcher et al., 2012; Vos et al., 2015) can now provide allele intensity information of thousands of single nucleotide polymorphisms (SNPs). After dosage calling is carried out (Schmitz Carley et al., 2017; Zych et al., 2019), these variants can be ultimately utilized in several applications such as linkage and quantitative trait locus (QTL) analyses (Hackett et al., 2014; Chen et al., 2018; Pereira et al., 2020b), genome-wide association studies (GWAS; Rosyara et al., 2016; Yuan et al., 2019), or genomic-assisted prediction (Sverrisdóttir et al., 2017; Enciso-Rodriguez et al., 2018). In the case of QTL identification of resistance to common scab in potato, only two studies have been carried out, one in a tetraploid population (227 F_1_ clones; Bradshaw et al., 2008), and another in a diploid population (49~91 F_2_ clones; Braun et al., 2017a). In addition, GWAS was performed in a tetraploid diversity panel (143 clones; Yuan et al., 2019).

In order to expand the understanding of the genetic control of common scab resistance in tetraploid potatoes, we evaluated a full-sib population with ~150 clones in three environments where common scab is of natural occurrence. A recently developed integrated genetic map based on a dosage-sensitive SNP chip array (Pereira et al., 2020b) was used for QTL mapping, and helped us to estimate haplotype-specific additive effects and to pinpoint candidate genes in the *S. tuberosum* genome that could potentially help breeders in deploying MAS for common scab in potato.

## 2 Material and Methods

### 2.1 F_1_ population and field trials

A mapping population named B2721, initially composed by 156 full-sibs, was derived from a cross between ‘Atlantic’ and B1829-5, and it was previously analyzed regarding its segregation to internal heat necrosis and several yield- and quality-related traits (McCord et al., 2011; Schumann et al., 2017; Pereira et al., 2020b). ‘Atlantic’ is a widely grown chipping variety in the USA, whereas B1829-5 is an advanced round white clone from the USDA-ARS Beltsville potato breeding program. Although both parents have shown susceptibility to common scab, B1829-5 was found to be less susceptible than ‘Atlantic’.

The B2721 was evaluated at three locations (hereafter also referred to as environments) in Becker, Minnesota in 2011 (MN11), in Pennsylvania Furnace, Pennsylvania in 2011 (PA11), and in Presque Isle, Maine in 2012 (ME12) in fields with a history of common scab pressure. In each location, the experimental design consisted of a randomized complete block with two replications, with four hills per plot. A total of 153 full-sibs were evaluated across locations, where MN11 included 146 full-sibs plus one check (B1829-5), PA11 included 151 full-sibs plus two checks (‘Atlantic’ and B1829-5), and ME12 included 139 full-sibs plus six checks (B1829-5, ‘Atlantic’, ‘Green Mountain’, ‘Ontario’, ‘Russet Burbank’ and ‘Superior’). All trials were carried out from early June (7~12) to late September (19~26) of their respective years. Standard crop management practices for the respective locations were followed.

All tubers were collected and visually assessed per plot for three traits of interest. First, the percentage of tubers with scab lesions in each plot (PS). Second, the tubers were rated for percentage of surface area covered by lesions (1 = <2% surface area; 2 = 2.1~5%; 3 = 5.1~10%; 4 = 10.1~25%; 5 = 25.1%~50%; 6 = >50%) following Merz scale (Merz, 2000), which was then converted to an area index (AI) as the sum of the individual tuber ratings of surface area infected divided by six times the number of tubers (Goth et al., 1993). Third, the tubers were also rated for type of lesion (0 = no lesions; 1 = superficial discrete; 2 = coalescing superficial; 3 = raised discrete; 4 = raised coalescing; 5 = pitted discrete and coalescing) following James (1971), which was then converted to a lesion index (LI) as the sum of the individual tuber ratings of lesion type divided by six times the number of tubers (Goth et al., 1993). The average number of tubers scored per plot was 30 in MN11, 24 in PA11, 21 in ME12.

### 2.2 Phenotypic analyses

Adjusted means were obtained based on a two-stage analysis approach using ASReml-R package v. 4.1.0 (Butler et al., 2018) and its restricted maximum likelihood (REML) estimation algorithm. In the first stage, phenotypic data was fitted for each separate environment using the model *y*_*ij*_ = *μ* + *b*_*j*_ + *g*_*i*_ + *ε*_*ij*_, where *y*_*ij*_ is the phenotypic value of individual *i* in block *j*, *μ* is the intercept, *b*_*j*_ is the fixed effect of block *j* (*j* = 1, …, *J*; *J* = 2), *g*_*i*_ is the fixed effect of individual *i* (*i* = 1, … , *n*; *n* = *n*_*g*_ + *n*_*c*_ with *n*_*g*_ = 139, 146 or 151 full-sibs, and *n*_*c*_ = 1, 2 or 6 checks depending on the environment), and *ε*_*ij*_ is the random effect of the residual error with *ε*_*ij*_ ~ *N*(0, *σ*^2^).

In the second stage, both adjusted means and weights derived from the diagonal of the variance-covariance inverse matrix from each first-stage model (Method 4 of Möhring and Piepho, 2009) were used to fit the model *μ*_*ik*_ *= ϕ* + *g*_*i*_ + *e*_*k*_ + *ge*_*ik*_ + *ϵ*_*ik*_, where *μ*_*ik*_ is the adjusted mean of individual *i* in environment *k* from the first-stage model, *ϕ* is the intercept, *g*_*i*_ is the fixed effect of individual *i*(*i* = 1, …, *n*), *e*_*k*_ is the random effect of environment *k* (*k* = 1, …, *K*; *K* = 3) with 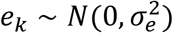, *ge*_*ik*_ is the random effect of genotype-by-environment interaction with 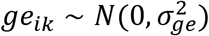, and *ϵ*_*ik*_ is the random effect of the residual error as a function of the weights from the first-stage model.

Approximate broad-sense heritability values were calculated using 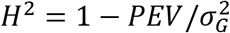 (Cullis et al., 2006; Isik et al., 2017, p. 223), where *PEV* is the best linear unbiased prediction error variance, and 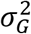 is the genetic variance associated with *g*_*i*_ when full-sib genotypes were treated as random in the two previous models, i.e. 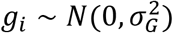. Pearson’s correlations were estimated between pairs of adjusted means derived from the first- and second-stage models using Hmisc R package v. 4.3-0 (Harrel Jr, 2019), correlograms were plotted using ggcorrplot2 R package v. 0.1.0 (Cai, 2019), and the remaining plots were obtained using ggplot2 R package v. 3.3.2 (Wickham, 2016).

### 2.3 Linkage mapping and QTL analyses

An integrated, fully phased genetic map was constructed using MAPpoly R package v. 0.1.0 (Mollinari et al., 2020) by Pereira et al. (2020b). This map is 1,630-centiMorgan (cM) long, comprises the 12 *S. tuberosum* base chromosomes, and contains 4,285 single nucleotide polymorphisms (SNPs) derived from the Illumina Infinium^®^ 8,303 Potato Array (Felcher et al., 2012). Genotype conditional probabilities were computed every cM using a hidden Markov model adapted to autopolyploids (Mollinari and Garcia, 2019), and ultimately employed in the QTL analyses.

For each trait, we used a random-effect multiple interval mapping (REMIM) model implemented in QTLpoly R package v. 0.2.1 (Pereira et al., 2020a) for QTL detection using the adjusted means derived from the phenotypic analyses. Variance components associated with putative QTL were tested using score statistics, whose *P*-values were compared to a genome-wide significance (*α*) assessed via score-based resampling method (Zou et al., 2004). In short, QTL were added (forward search) to a random-effect model using a more relaxed genome-wide significance level (*α* = 0.20). Then, QTL already in the model were re-evaluated under a more stringent significance level (*α* = 0.05) and excluded (backward elimination) if not significant. These steps were repeated under the more stringent significance level (*α* = 0.05), and the forward-backward algorithm was stopped once no more QTL were either added to or excluded from the model. Putative QTL were tested every cM position along the B2721 genetic map, and a 20-cM window on each side of QTL already in the model was avoided when searching for another QTL. The genotypic values derived from the final QTL model were used to compute additive allele effects (Pereira et al., 2020a).

## 3 Results

The reaction to common scab in the B2721 mapping population was evaluated across three environments, and the raw phenotypic data, as well as the adjusted means and weights, are made available in the Supplementary File S1.

All three evaluated traits showed broad phenotypic value ranges, with several full-sib clones showing transgressive segregation, i.e. being more resistant or more susceptible when compared to parental means (Table 1). Although similar ranges were observed for all evaluated traits across environments, reaction to common scab in PA11 and ME12 were skewed towards more severe phenotypes, whereas MN11 behaved in the opposite direction, hence towards resistance (Figure 1a-c). In fact, the mapping population mean of percentage of scabby tubers (PS) was only 18.88% in MN11, but 91.64% and 81.15% in PA11 and ME12, respectively (Table 1). Similar trends were observed for area index (AI), with 0.07 in MN11, but 0.27 and 0.31 in PA11 and ME12, and lesion index (LI), with 0.13 in MN11, but 0.60 and 0.64 in PA11 and ME12. The common check, B1829-5, also performed very inconsistently across locations, showing 95.22% and 85.00% of scabby tubers in PA11 and ME12, respectively, but only 3.85% in MN11. Variation for these traits in MN11 could not be attributed to genetics, as evidenced by their null heritability estimates. In PA11 and ME12, the heritability values were 0.53 and 0.72 for PS, 0.48 and 0.70 for AI, and 0.78 and 0.79 for LI, respectively. For the joint model, the respective heritability values for PS, AI and LI were 0.48, 0.44 and 0.67 (Table 1).

**Table 1.** Summary of percentage of scabby tubers (PS), area index (AI) and lesion index (LI) of common scab reaction in the B2721 potato mapping population based on separate (MN11, PA11, ME12) and joint analyses.

| Trait | Location | Means | | | F<sub>1</sub> range | $\sigma_G^2$<sup>b</sup> | $H^2$<sup>c</sup> |
| --- | --- | --- | --- | --- | --- | --- | --- |
|  |  | ‘Atlantic’ | B1829-5 | F<sub>1</sub> |  |  |  |
| PS | Joint | 72.32 | 65.12 | 63.68 | 16.13-78.40 | 5.61E+01 | 0.485 |
|  | MN11 | NA<sup>a</sup> | 3.85 | 18.88 | 0.00-86.93 | 4.32E-05 | 0.000 |
|  | PA11 | 100.00 | 95.22 | 91.64 | 53.57-100.00 | 4.90E+01 | 0.531 |
|  | ME12 | NA | 85.00 | 81.15 | 0.00-100.00 | 2.65E+02 | 0.724 |
| AI | Joint | 0.29 | 0.24 | 0.22 | 0.02-0.44 | 2.64E-03 | 0.437 |
|  | MN11 | NA | 0.01 | 0.07 | 0.00-0.49 | 8.11E-10 | 0.000 |
|  | PA11 | 0.33 | 0.28 | 0.27 | 0.09-0.50 | 3.19E-03 | 0.481 |
|  | ME12 | NA | 0.40 | 0.31 | 0.00-0.64 | 1.48E-02 | 0.702 |
| LI | Joint | 0.64 | 0.54 | 0.45 | 0.02-0.77 | 1.87E-02 | 0.668 |
|  | MN11 | NA | 0.01 | 0.13 | 0.00-0.74 | 2.60E-09 | 0.000 |
|  | PA11 | 0.79 | 0.80 | 0.60 | 0.11-1.00 | 3.67E-02 | 0.777 |
|  | ME12 | NA | 0.72 | 0.64 | 0.00-0.98 | 3.70E-02 | 0.791 |

<sup>a</sup>NA = not available <sup>b</sup> $\sigma_G^2$ = genetic variance <sup>c</sup> $H^2$ = broad-sense heritability by Cullis et al. (2006)

**Figure 1.**
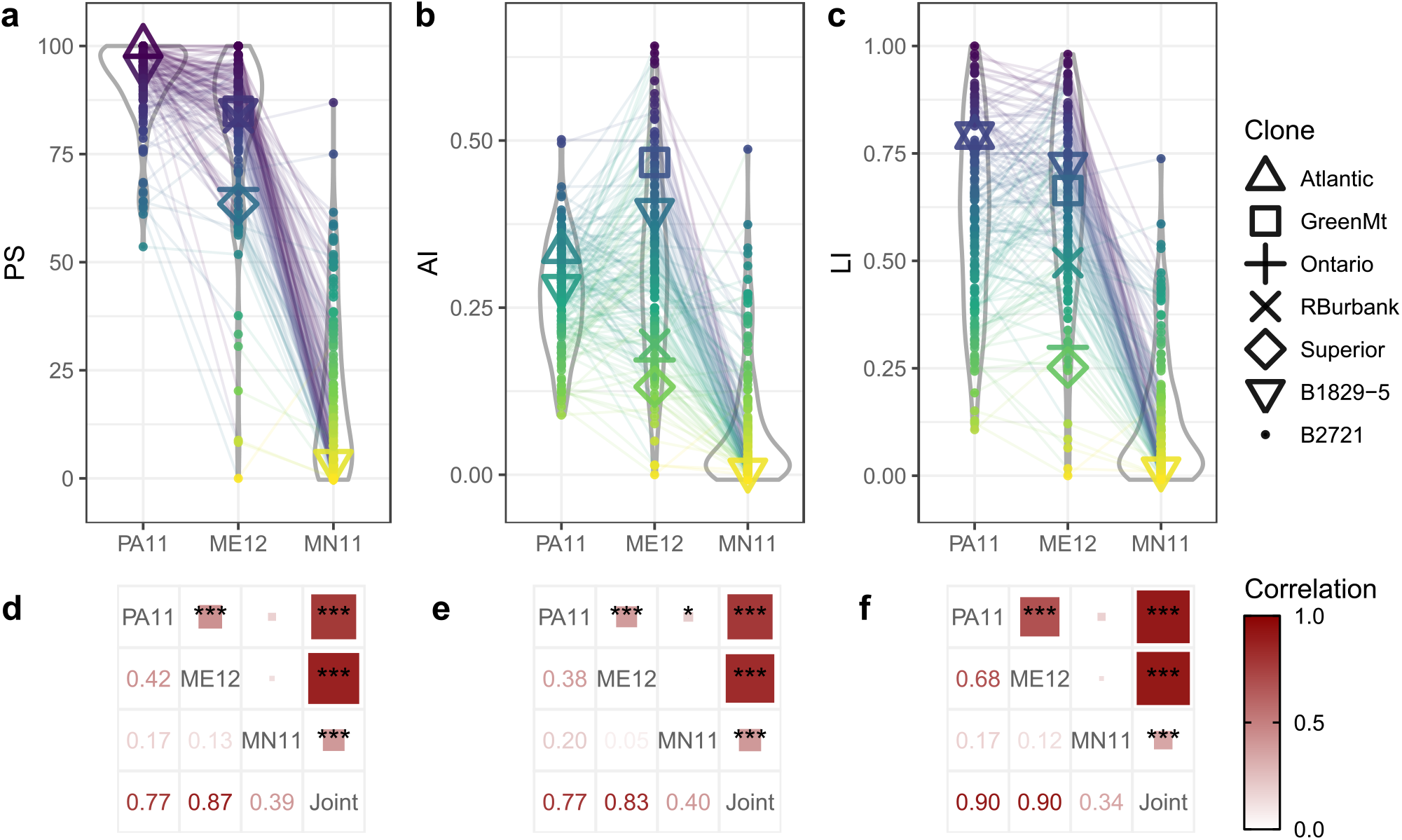
Violin (a-c) and correlogram (d-f) plots for percentage of scabby tubers (PS; a and d), area index (AI; b and e) and lesion index (LI; c and f) of common scab reaction in the B2721 potato mapping population evaluated across three environments (MN11, PA11, ME12). Symbols represent different checks. Lines connect the same B2721 clone in consecutive environments. Correlograms show Pearson’s correlation values (\**P* < 0.05, \*\*\**P* < 0.001) between separately and jointly adjusted means.

The correlation among separately adjusted means for different environments can be observed in Figure 1d-f. Between PA11 and ME12 means, the correlations were positive and moderate for PS (0.42), AI (0.38) and LI (0.68), but low between these environments and MN11 (0.05~0.20). The lack of consistency in the individual ranking across environments can be visualized in Figure 1a-c, especially in relation to MN11, implying strong genotype-by-environment interaction. The correlation estimates between the jointly adjusted means with the PA11 and ME12 means were high (0.77~0.90), and with MN11 were moderate (0.34~0.40). As these three environments were under natural common scab pressure, meaning that no inoculum was artificially applied to the clones, we believe that in MN11 certain environmental conditions, including availability of less pathogenic *Streptomyces* spp. or strains, did not allow the disease to progress as much as in PA11 and ME12. Although MN11 had little to offer for genetic analysis purposes, the adjusted means derived from this environment were carried along with the analysis as a sort of negative control for QTL mapping.

We also compared the correlation among traits using their jointly adjusted means (Figure 2). We observed that the traits were highly, positively correlated (0.76~0.83). This is likely related to the progression of the disease, such that in a susceptible genotype, more scabby tubers resulted in broader areas covered by scab-like lesions, which in turn appeared to be more severe. Despite the high correlation, we were able to identify QTL only for LI. QTL mapping models were fitted using separately and jointly adjusted means derived from the phenotypic analyses. The QTL for LI was co-localized on linkage group 3 at 99 cM for PA11, ME12 and jointly adjusted means, but no QTL was found for MN11 (Figure 3a). This QTL explained as much as 30.1% of the trait variation in PA11. For ME12 and jointly adjusted means, the QTL heritability was 22.5% and 25.4%, respectively (Table 2). The significance of the score statistics (*P*-values) ranged from 4.17E–05 (ME12) to 1.05E– 06 (PA11).

**Figure 2.**
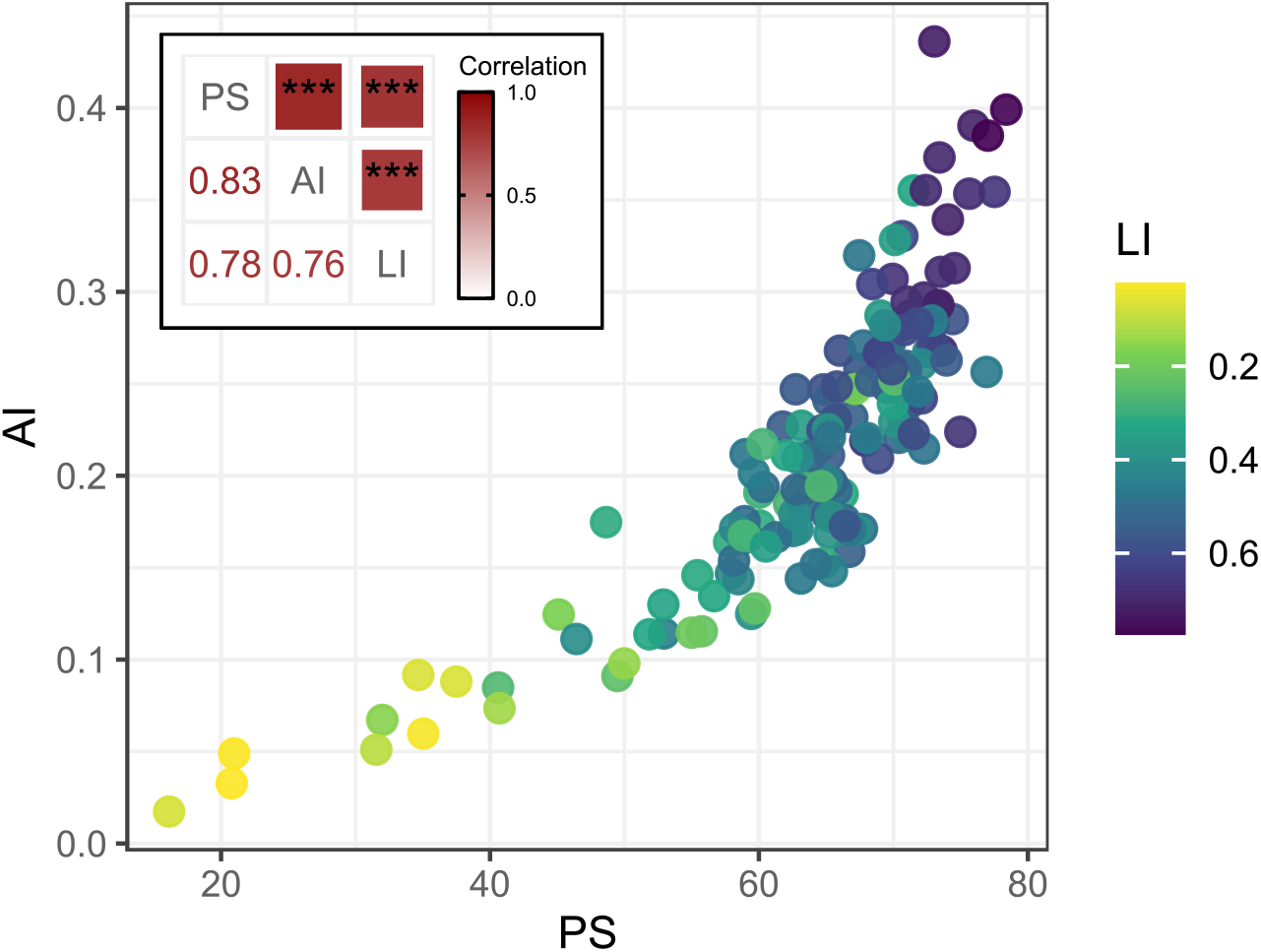
Scatterplot of jointly adjusted means for percentage of scabby tubers (PS), area index (AI) and lesion index (LI) of common scab reaction in the B2721 potato mapping population evaluated across three environments. Correlogram (top-left) shows Pearson’s correlation values (\*\*\**P* < 0.001).

**Figure 3.**
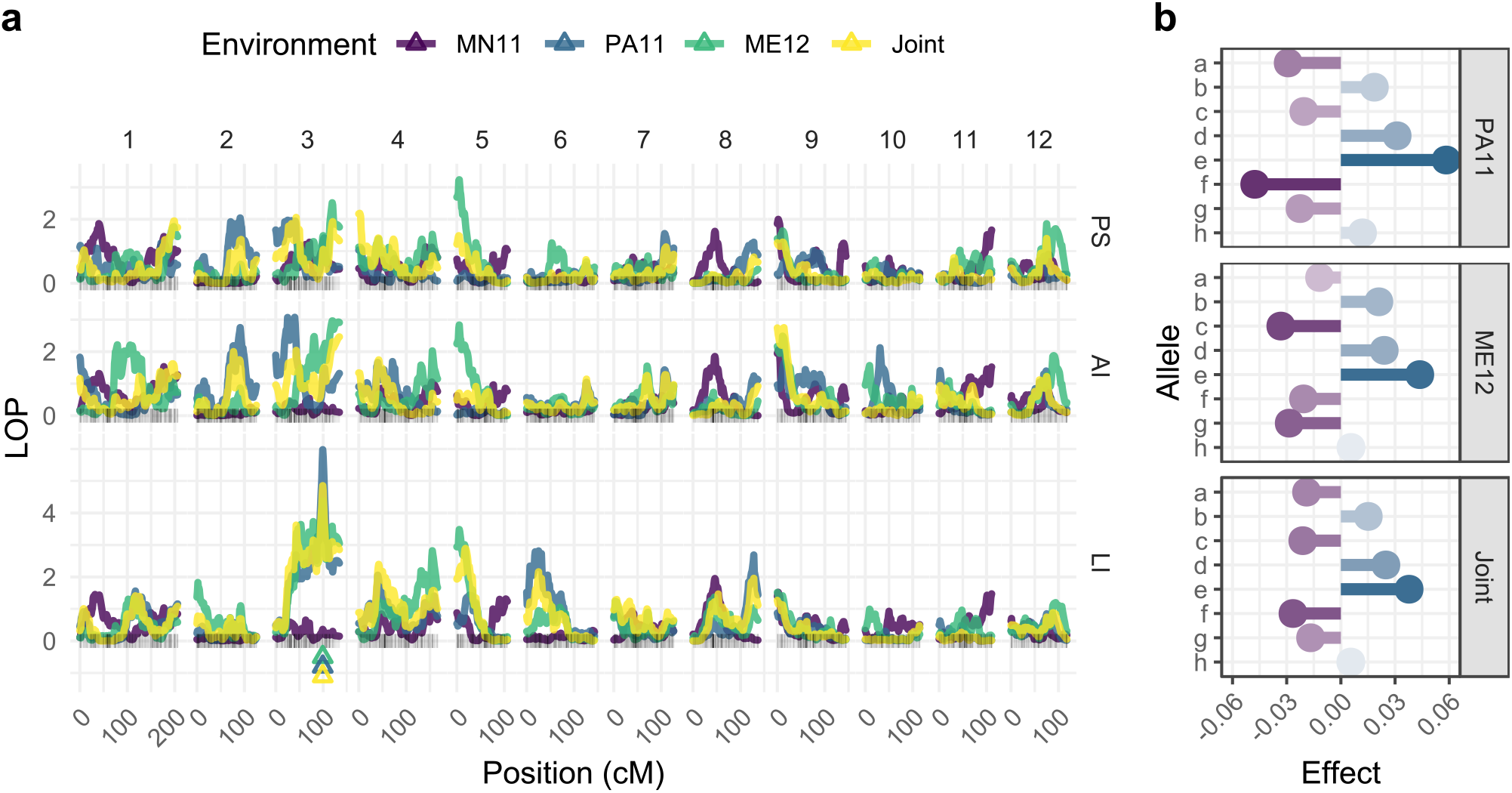
(a) QTL profiles for percentage of scabby tubers (PS), area index (AI) and lesion index (LI) of common scab reaction in the B2721 potato mapping population evaluated across three environments (MN11, PA11, ME12). “Joint” refers to the QTL analysis using the jointly adjusted means, triangles represent the QTL peak, and *LOP* = − log_10_(*P*). (b) Additive effects of the QTL on linkage group 3 at 99 cM for LI in PA11, ME12 and Joint analyses. Parental alleles (haplotypes): ‘Atlantic’ = *a*, *b*, *c*, *d*, and B1829-5 = *e*, *f*, *g*, *h*.

**Table 2.** QTL detected for lesion index (LI) of common scab reaction in the B2721 potato mapping population based on separate (PA11, ME12) and joint analyses.

| Environment | LG<sup>a</sup> | Position (cM) | SI<sup>b</sup> (cM) | Score | P-value | Intercept | $\sigma_{QTL}^2$<sup>c</sup> | $h_{QTL}^2$<sup>d</sup> |
| --- | --- | --- | --- | --- | --- | --- | --- | --- |
| PA11 | 3 | 99 | 96-103 | 195.32 | 1.05E-06 | 0.587 | 0.0162 | 0.301 |
| ME12 | 3 | 99 | 43-134 | 131.90 | 4.17E-05 | 0.626 | 0.0110 | 0.225 |
| Joint | 3 | 99 | 43-103 | 161.17 | 1.40E-05 | 0.446 | 0.0078 | 0.254 |

<sup>a</sup>LG = linkage group <sup>b</sup>SI = ~95% support interval <sup>c</sup> $\sigma_{QTL}^2$ = variance component associated with the QTL <sup>d</sup> $h_{QTL}^2$ = QTL heritability

The QTL allele contributions for LI were relatively consistent among PA11, ME12 and joint analyses (Figure 3b). The parent B1829-5 showed more pronounced additive effects, with haplotype *e* contributing to increasing (+0.0583) and haplotype *f* contributing to decreasing (–0.0476) the mean (0.587) LI in PA11. In this case, in addition to haplotype *f*, one would have to select haplotype *g* from B1829-5, and haplotypes *a* and *c* from ‘Atlantic’ in order to potentially drive LI down.

The SNP markers at ~95% support interval boundaries of the QTL in PA11 (96~103 cM) were solcap_snp_c2_1830 (ST4.03ch3:51731168) and solcap_snp_c1_7076 (ST4.03ch3:54368685). This region spanned 2,637,517 bp (4.23%) of the chromosome 3 of the S. tuberosum v. 4.03 reference genome (Sharma et al., 2013), and contained 263 genes (Supplementary File S2). The closest markers on the left and on the right of the QTL peak were solcap_snp_c2_57263 (ST4.03ch3:53439319) and solcap_snp_c1_5812 (ST04.03ch3:53665337), respectively, and spanned 226,018 bp; this region contained 23 genes.

## 4 Discussion

While both B2721 parents and most of the full-sibs are highly susceptible to common scab for two environments, namely PA11 and ME12, the absence of more severe symptoms in MN11 can be attributed to a strong genotype-by-environment interaction. The variation observed in this environment is hence due to non-genetic, environmental effects. For all traits, the lack of correlation between MN11 and the other two environments confirms such a divergent pattern. This also explains how there was no evidence for QTL in the same region as for PA11 and ME12, where common scab was relatively more severe. That is, in order to map QTL for resistance to a disease, the environment pathogen pressure should be such that the genotypes can express their genetic merit and be phenotypically evaluated. In the case of potato common scab, the variation present within pathogenic *Streptomyces* spp., and other soil variables, such as moisture content and pH, are known to influence the severity of scab (Braun et al., 2017b). The distribution of different *Streptomyces* spp. isolates predominating in different parts of the USA (Wanner, 2009) could partially explain the lack of agreement between MN and either ME or PA.

Bradshaw et al. (2008), working with 227 full-sib progenies, encountered similar issues when only one out of three environments showed scorable severity for common scab, for which heritability was 0.66. While studying 23 tetraploid potatoes, Haynes et al. (1997) found higher broad-sense heritability values for AI (0.89) and LI (0.93), where a rather low genotype-by-environment interaction was observed. For 370 clones evaluated over nine years, Enciso-Rodriguez et al. (2018) found low genotype-by-year interaction and a genomic heritability estimate of 0.45 for common scab scoring. In another study involving diploid potatoes, where lesion type and percentage of surface area was scored for common scab over three years, Braun et al. (2017b) found heritability values ranging from 0.48 to 0.79. Finally, in a diversity panel with 148 clones, heritability was estimated as 0.81 for a three-year evaluation data (Yuan et al., 2019). Therefore, except for the null heritabilities resulting from MN11, our estimates (0.44~0.79) were in relative agreement with those found in literature.

Some genomic regions have been found to be associated to common scab resistance, but none on chromosome 3 as identified in the present study. Previous QTL mapping studies in tetraploid (Bradshaw et al., 2008) or diploid (Braun et al., 2017a) populations found two (on homology groups II and IV) or one QTL (on chromosome 11), respectively. Using GWAS, Yuan et al. (2019) found associations on chromosomes 2, 4, and 12. Modernly, molecular breeding has been taking advantage of the high-density markers covering the whole genome to perform selection based on genomic estimated breeding values. For potato common scab, such genomic-assisted prediction models have shown prediction accuracies as high as 0.278, and a SNP with major effect on chromosome 9 (Enciso-Rodriguez et al., 2018). Haplotypic and QTL information, such as that provided here, can be used to leverage genomic-assisted prediction models in order to deliver higher predictive abilities, notably for less complex traits (Gemenet et al., 2020).

At least, three out of the 23 genes located within the QTL marker interval were previously implicated in plant responses to biotic stresses in signaling pathways or hypersensitive responses, namely MYB transcription factor (PGSC0003DMT400063172, ~53.60 Mbp) (Ambawat et al., 2013), calcium-dependent protein kinase 1 (CDPK1; PGSC0003DMT400034291, ~53.43 Mbp) (Lee and Rudd, 2002), and ubiquitin-protein ligase (PGSC0003DMT400092368, ~53.59 Mbp) (Craig et al., 2009; Duplan and Rivas, 2014). In addition, there were several other genes among the remaining 240 transcripts retrieved from the ~95% QTL support interval that were known for encoding proteins involved in plant defense, such as receptor-like kinase (PGSC0003DMT400046807, ~52.47 Mbp; PGSC0003DMT400046778, ~52.57 Mbp) (Nazarian-Firouzabadi et al., 2019), proteins containing nucleotide biding-ARC domain (PGSC0003DMT400087730, ~52.37 Mbp) and leucine-rich repeat (LRR; PGSC0003DMT400046655, ~52.18 Mbp) (Takken et al., 2006), and transcription factors such as NAC (PGSC0003DMT400097372, ~54.15 Mbp; PGSC0003DMT400096000, ~54.17 Mbp) (Nuruzzaman et al., 2013) and WRKY (PGSC0003DMT400046570, ~53.04) (Bhattarai et al., 2010; Enciso-Rodriguez et al., 2018).

Although the list of candidate genes is still hypothetical, several transcripts encoding related protein isoforms (e.g. MYB, WRKY, LRR receptor-like serine/threonine-protein kinase) were found to be differentially expressed between the resistant ‘Hindenburg’ and susceptible ‘Green Mountain’ cultivars inoculated with *S. scabies* (Fofana et al., 2020). Here, we detected a single QTL consistently in two out of three environments, that explains up to 30% of the variation for lesion index. In order to effectively apply genes and haplotypes in breeding, the QTL identified in this study needs to be further investigated in a more diverse genetic background. If an oligogenic-like inheritance confirms and upon QTL validation, specific markers to retrieve the haplotypes conferring resistance to common scab can be screened in breeding populations to perform early selection (Bradshaw, 2017). Progress of MAS in potato is relatively slow when compared to diploid, inbred species, and needs to take into consideration several aspects in addition to the genetic architecture of a trait, such as polyploidy, high heterozygosity, and clonal propagation (Slater et al., 2014; Bethke et al., 2019).

## 6 Author Contributions

KH designed the experiments and supervised the project. KH, XQ and CT performed field experiments and collected phenotypic data. CY provided genotypic data. GSP analyzed data and drafted the manuscript. MM and ZBZ supervised data analyses. All authors reviewed and approved the manuscript.

## 8 Data Availability Statement

The phenotypic data analyzed for this study can be found as Supplementary Material. The genotypic data and linkage map information are available at https://github.com/mmollina/B2721_map (Pereira et al., 2020b). MAPpoly (https://github.com/mmollina/mappoly) and QTLpoly (https://github.com/guilherme-pereira/QTLpoly) source codes are available at their respective GitHub pages.

## 9 Supplementary Material

Supplementary File S1. Raw phenotypic data, adjusted means and weights from phenotypic analyses. (XLSX)

Supplementary File S2. List of annotated genes within the ~95% support interval for the QTL for lesion index (LI) in PA11. (XLSX)

